# Inefficient transmission of African swine fever virus to sentinel pigs from environmental contamination under experimental conditions

**DOI:** 10.1101/2023.09.28.559902

**Authors:** Ann Sofie Olesen, Louise Lohse, Francesc Accensi, Hannah Goldswain, Graham J. Belsham, Anette Bøtner, Christopher L. Netherton, Linda K. Dixon, Raquel Portugal

## Abstract

Knowledge about African swine fever virus (ASFV) transmission and its survival in the environment is mandatory to develop rational control strategies and combat this serious disease in pigs. In this study, the risk that environmental contamination poses for infection of naïve pigs was investigated. Naïve pigs were introduced as sentinels into contaminated pens either on the same day or up to three days after ASFV-infected pigs were removed. Three experiments were carried out in which four to six pigs per pen were inoculated with virulent ASFV isolates OURT88/1 (genotype I), Georgia 2007/1 or POL/2015/Podlaskie (genotype II), respectively. The majority of the inoculated pigs developed acute disease but with no evident haemorrhagic lesions or haemorrhagic diarrhoea and were culled at the predefined humane endpoint. The levels of ASFV DNA detected in the blood of the infected animals reached 10^7-9^ genome copies/ml before euthanasia. Environmental swabs were taken from different surfaces in the animal rooms, as well as from faeces and urine, close to the time of introduction of the naïve animals. Relatively low quantities of virus DNA were detected in the environmental samples, in the order of 10^3-7^ genome copies. Neither clinical signs nor virus genomes were detected in the blood of any of the sentinel pigs over a period of two to three weeks after exposure, indicating that transmission from the ASFV-contaminated environment did not occur. Interestingly, viral DNA was detected in nasal and oral swabs from some of the sentinel animals at early days of exposure (ranging between 10^3.7-5.8^ genome copies), though none of them developed ASF. The results indicate a relatively low risk of ASFV transmission from a contaminated environment in the absence of blood from infected animals.

## 1. Introduction

African swine fever (ASF) is a severe haemorrhagic disease with a high case fatality rate in domestic pigs and wild boar. It is caused by African swine fever virus (ASFV), a large, cytoplasmic, double stranded DNA virus that is the only member of the *Asfarviridae* family. Safe and efficient commercial vaccines are not yet available to aid disease control. A long-established wildlife reservoir of ASFV is present in East Africa in warthogs and soft ticks from the genus *Ornithodoros* that inhabit their burrows. These and other wild suids in Africa, including bush pigs, show no disease and develop a low transient viremia but animals can remain persistently infected for long periods (Dixon, Stahl et al. 2020, Penrith and Kivaria 2022). In contrast, infected domestic pigs and wild boar develop high titres of virus in blood and direct transmission occurs readily between them (Sauter-Louis, Conraths et al. 2021). The large numbers of wild boar in many European countries provide a wildlife reservoir for infection of domestic pigs.

Indirect transmission by various mechanisms is also recognised as an important transmission route (Olesen, Belsham et al. 2020, Nielsen, Alvarez et al. 2021). Ingestion of pork products containing infectious virus is important and is often the route for long distance virus spread. Fomites such as clothing or transport trucks can also be a source of infection although this route was estimated to be less important. Contaminated feed or water supplies as well as wild boar carcasses can also provide sources of infection for spread of the virus into wild boar and spill over into domestic pigs (Bergmann, Schulz et al. 2021). Mechanical transmission by biting flies has been suggested to play a role in virus transmission but has been little studied. Two studies showed that ASFV survived for 48 hours in stable flies fed on infected blood with high viremia (10^8^ HAD_50_/ml) (Mellor, Kitching et al. 1987) or 12h after feeding on blood with lower viremia (5 x 10^5^ TCID_50_/ml) (Olesen, Hansen et al. 2018). In the study by Mellor et al. (Mellor, Kitching et al. 1987), ASFV was transmitted to pigs by the biting flies that had been blood fed 1h and 24h before feeding on the animals. It has also been shown that ingestion of stable flies fed on ASFV infected blood also resulted in infection of pigs (Olesen, Lohse et al. 2018). Furthermore, in an ASF outbreak area, hematophagous insects carrying blood meals including ASFV DNA were captured on the windows of a high biosecurity pig farm that was free of ASF, hence indicating a potential risk for introduction of ASFV (Stelder, Olesen et al. 2023).

Aerosol transmission of ASFV has been detected over short distances within buildings but wider dispersion by aerosol is not thought to occur (Wilkinson, Donaldson et al. 1977, Olesen, Lohse et al. 2017). As a large DNA virus, ASFV is physically very stable and can survive for extended periods in contaminated materials posing additional problems for control. A recent EFSA Scientific Opinion reviewed literature on survival of ASFV in different matrices and estimated the risk these posed for virus transmission in different scenarios (Nielsen, Alvarez et al. 2021). Very high levels of virus are present in the blood of pigs showing clinical signs of acute ASF (up to 10^8-9^ TCID_50_ or HAD_50_/ml). A very early study showed that blood collected after death with ASF and stored at room temperature in the dark for 140 days was still infectious as shown by inoculation of another pig (Montgomery 1921). ASFV was also observed to survive in chilled blood for an extended period of time, some 525 days (Plowright and Parker 1967). Thus, materials contaminated with infected blood pose a high risk for spread of virus. Although much lower levels of virus are found in excretions, ASFV can also survive in these for several days. In faeces, with an initial titre of 10^4.83^ TCID_50_/g, collected from animals showing acute disease, ASFV survived for up to 8.5 days when the faeces were stored at 4°C, for 5 days at 21°C and 3 days at 37°C (Davies, Goatley et al. 2017). In an early study, virulent virus was still present in faeces kept at room temperature in the dark for up to 11 days, with these faeces inducing ASF and death in seven days after being fed to a susceptible pig (Montgomery 1921). This study also found that virus in urine collected after death survived at room temperature for at least 2 days, and caused ASF after being fed to a susceptible pig. However, after storage for longer than 2 days, the urine was less likely to cause infection. More recently, it was observed that ASFV in the urine from infected animals, with initial titres of 10^2.2-3.8^ TCID_50_/ml, could survive, on average, for 15 days when chilled, for 5 days at room temperature or for 3 days when incubated at 37°C (Davies, Goatley et al. 2017). In agreement with these results, a recent study did not detect infectious virus in faeces and urine from ASFV-infected pigs and wild boar after one week of storage at room temperature (Fischer, Huhr et al. 2020).

Water troughs are shared by the animals in a pig pen and hence if water becomes contaminated it may also spread infection, especially since ASFV seems to be highly stable in water. Experimentally contaminated water, with an initial titre of 10^6.5^ HAD_50_/ml stored frozen (−16 to −20°C) or chilled (4– 6°C), contained viable ASFV for at least 60 days, and when stored at room temperature (22–25°C) was infectious for 50 days (Sindryakova 2016).

No data were found estimating ASFV survival on straw, a commonly used bedding material in pig farms (Nielsen, Alvarez et al. 2021). However, it would be expected that straw in housing with infected pigs may be contaminated with faeces and urine as well as blood and hence constitute an important source of transmission.

The infectious dose of ASFV varies according to the route of infection. In a recent study the minimum infectious dose of virulent ASFV genotype II in liquid was estimated to be 10^0^ TCID_50_, compared with 10^4^ TCID_50_ in feed (median infectious dose was 10^1.0^ TCID_50_ for liquid and 10^6.8^ TCID_50_ for feed) (Niederwerder, Stoian et al. 2019). This is a large difference, for which the basis is not known. It has been suggested that liquids provide a suitable substrate for contact between the virus and the tonsils. In an earlier study, the intranasal/oral infectious dose_50_ (ID_50_), and the intravenous/intramuscular (IV/IM) ID_50_ of a moderately virulent isolate of ASFV were determined to be 18,500 and 0.13 HAD_50_, respectively, and a highly virulent isolate required approximately 10-fold more virus to cause infection by the intranasal/oral route (McVicar 1984). Although these results vary in their estimates of infectious dose, both confirm that ASFV can readily be transmitted by the oral-nasal route. Since infectious virus can survive for several days, at the range of temperatures where pigs are reared, blood and secretions may pose a risk for transmission.

Although progress has been made in understanding mechanisms and risks posed by different indirect routes of ASFV transmission, gaps in knowledge remain. In early studies, transmission to pigs from an environment contaminated with ASFV was observed when a contaminated pen had been left empty for three days, but not for five days (Montgomery 1921). In more recent experiments (Olesen, Lohse et al. 2018; Olesen, 2019), it was shown that pigs that were introduced into the contaminated environment at 1 day after removal of ASFV-infected animals, developed clinical disease. However, pigs introduced into the contaminated pens after 3, 5 or 7 days did not develop signs of ASF. The results suggested a relatively narrow window of time for transmission, but further studies were needed to confirm this.

In the current study we investigated the potential role of environmental contamination in pig housing for transmission of ASFV. Three experiments were carried out in which naïve pigs were introduced into pens that had recently housed pigs showing acute disease after inoculation with virulent isolates of ASFV: OURT88/1 (genotype I), Georgia 2007/1 or POL/2015/Podlaskie (genotype II). The naïve pigs were introduced on the same day or 1-3 days after the infected pigs were removed and the levels of viral DNA in different surface swabs and excretions were evaluated at different days before and during exposure. None of the introduced pigs became infected suggesting that the risk of transmission from environmental contamination is low, although we detected relatively low levels of virus genome in environmental samples collected from rooms that had housed the infected pigs and also in some oral/nasal swabs from the introduced animals.

## 2. Material and Methods

### 2.1. Virus Isolates and Cell Culture

The OURT88/1 (genotype I) and Georgia 2007/1 (genotype II) virulent isolates of ASFV have been described previously (Boinas, Hutchings et al. 2004) (Rowlands, Michaud et al. 2008). Virus stocks were prepared by infection of primary porcine bone marrow cells and titrated by limiting dilution in porcine bone marrow cells seeded in 96 well plates using a hemadsorption (HAD) assay and are expressed as HAD_50_ /ml as described previously (Goatley, Reis et al. 2020). The ASFV POL/2015/Podlaskie (genotype II) was isolated as previously described (Olesen, Lohse et al. 2017). The virus was prepared by infection of porcine pulmonary alveolar macrophages (PPAM) and titrated in PPAM seeded in 96 well plates using an immunoperoxidase monolayer assay (IPMA) as described previously (Botner, Nielsen et al. 1994, Olesen, Lohse et al. 2017) with titres presented as TCID_50_/ml (REED and MUENCH 1938).

### 2.2. Animal housing and Ethical approval

Animal experiments 1 and 2 were carried out at The Pirbright Institute under license 7088520 issued by the UK Home Office under the Animals (Scientific Procedures) Act (1986) (ASPA) and were approved by the Animal Welfare and Ethical Review Board (AWERB). The animals were housed in the SAPO4 high containment large animal unit at The Pirbright Institute in accordance with the Code of Practice for the Housing and Care of Animals Bred, Supplied or Used for Scientific Purposes. Bedding and species-specific enrichment were provided throughout the study to ensure high standards of welfare. Clinical scoring was carried out daily (King, Chapman et al. 2011) and pigs that reached the scientific or moderate severity humane endpoint, as defined in the project license, were euthanized by an overdose of anaesthetic.

Animal experiment 3 was performed in BSL3 facilities at the *Centre de Recerca en Sanitat Animal* (IRTA-CReSA, Barcelona, Spain). The experiment was conducted in accordance with EU legislation on animal experimentation (EU Directive 2010/63/EU). A commercial diet for weaned pigs and water were provided *ad libitum*Rectal temperatures and clinical signs were recorded for each pig on a daily basis. A total clinical score was calculated per day based using a previously described system (Olesen, Kodama et al. 2021; Olesen, Lazov et al. 2023). Pigs were euthanized, after reaching the humane endpoints set in the study, by intravascular injection of Pentobarbital following deep anesthesia.

### 2.3. Animal experiments design

For experiments 1 and 2, female Landrace × Large white × Hampshire pigs were obtained from a high health status farm in the UK and after a seven-day settling-in period were challenged intramuscularly with 10,000 HAD_50_ infectious units of virulent ASFV. In experiment 1, five animals were inoculated with OURT88/1 isolate. In experiment 2, six animals were inoculated, three with OURT88/1 and the other three with Georgia 2007/1, all kept in the same pen. The infected animals were euthanized five days after infection when reaching the humane endpoint. The premises were minimally cleaned between days 3-5 of infection (removal of gross faeces contamination and any blood post-sampling, as well as ensuring the pigs had a clean area to eat) and after that were left completely uncleaned until two sentinel pigs were introduced into the room. In Experiment 1, sentinels were introduced to the premises one day after removal of the infected animals and, in Experiment 2, they were introduced on the same day. Minimal cleaning was then restarted the following day and the sentinel animals were monitored for clinical signs over a period of 14 days. Figure 1 shows the design of experiment 1 including days on which samples were collected. During the experiments, temperature and relative humidity of the premises housing the animals were recorded daily and were 18.4-19.1⁰C and 45-53.2% respectively. Air exchange rates in the animal housing rooms were approximately 13 per hour.

**Figure 1:**
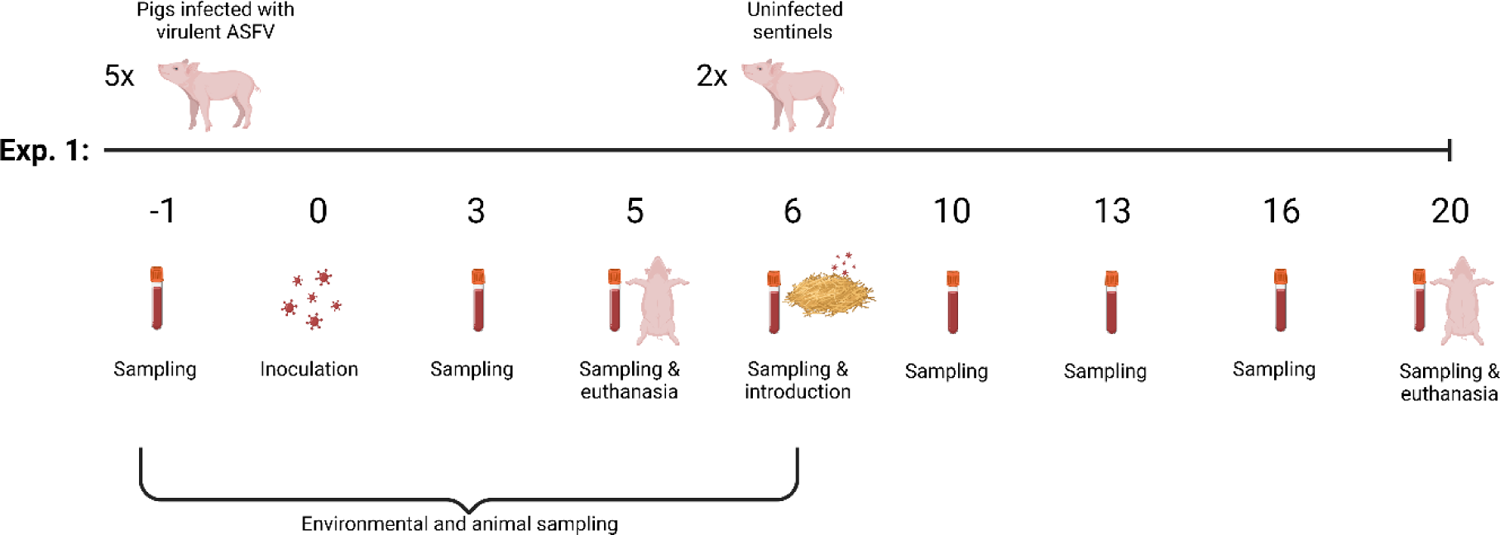
Plan of animal experiment 1 involving animal and environmental sampling and sentinel monitoring for ASF transmission. Two sentinel pigs were introduced into the premises one day after euthanasia of infected controls. Numbers below the line represent days of the experiment with reference to the inoculation day (0).

For experiment 3, 24 male Landrace X Large White pigs were obtained from a conventional Spanish swine herd. After an acclimatization period of one week, twelve pigs (pigs 1-12) were challenged intranasally with 10,000 TCID_50_ as already described (Olesen, Kodama et al. 2021). The infected animals were euthanized six days after inoculation when the humane endpoint was reached. The twelve inoculated pigs were housed in three separate high containment units (termed boxes 4, 5 and 6): Pigs 1-4 in box 4, pigs 5-8 in box 5 and pigs 9-12 in box 6 (Figure 2). The three pens were identically designed with slatted (2/3) and solid (1/3) flooring. The three boxes had a room volume of 70 m^3^, an average temperature of 22°C (±0.19°C) and 11-16 air renewals per hour. No cleaning was performed during the course of the experiment. Enrichment materials (rope, toys) were available within all pens. Following euthanasia of the twelve inoculated pigs, all material, including feed, faeces and toys, remained in the pens. In order to avoid excessive drying, the pen floors were covered with plastic (approximately 52 cm above the floor). Under these conditions, the three pens within boxes 4, 5 and 6 were left empty for 1-, 2- or 3-days following removal of the ASFV-infected pigs. Subsequently, twelve sentinel pigs (numbered 13-24), were introduced into the contaminated pen environments in the following order: pigs 13-16 were introduced into box 6 at 1 day, pigs 17-20 into box 5 at 2 days, and pigs 21-24 into box 4 at 3 days post-euthanasia of the ASFV-infected pigs, respectively (Figure 2).

**Figure 2:**
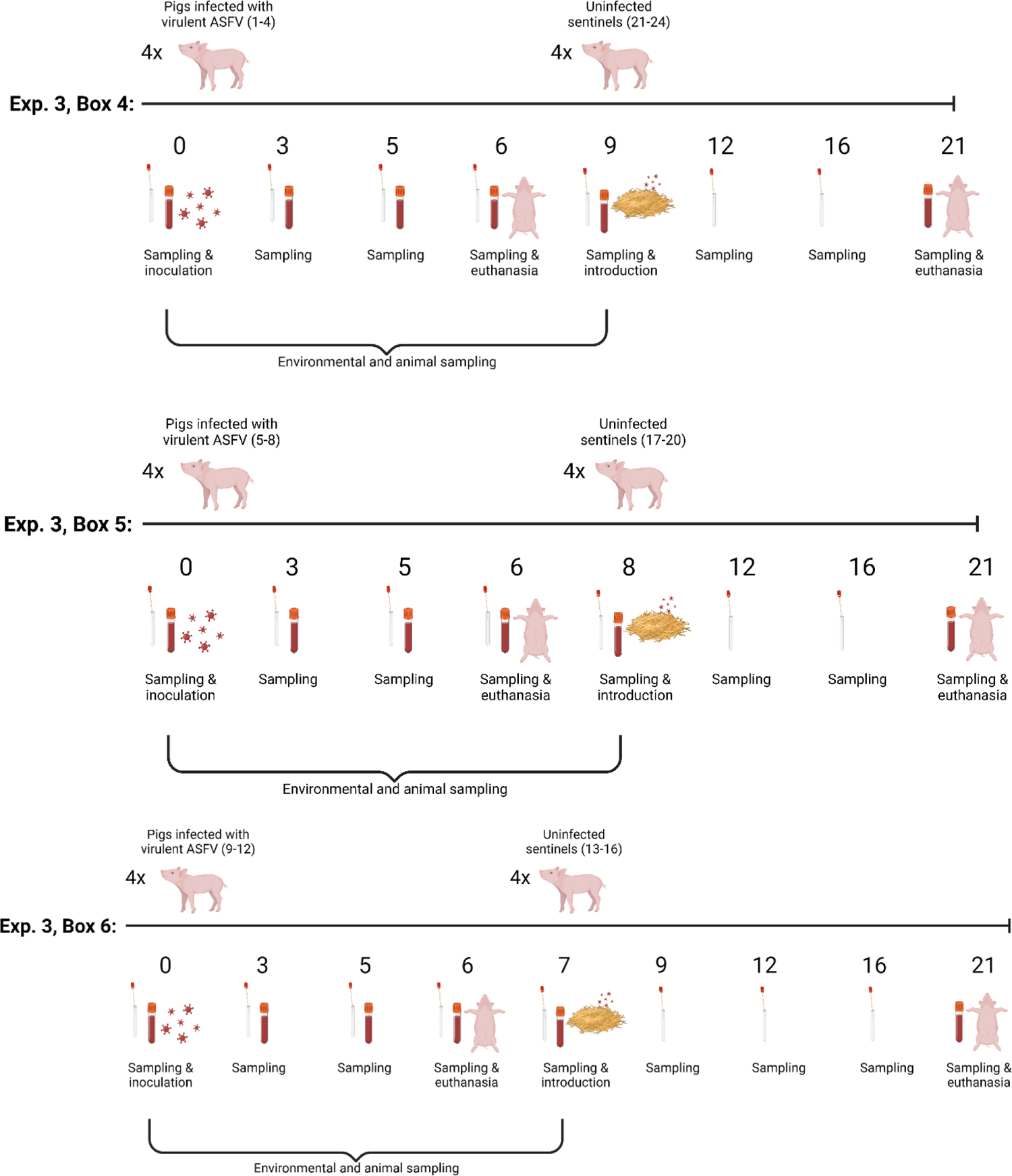
Overview of study design in boxes 4-6 in experiment 3. Numbers below the line represent days of the experiment relative to the inoculation day (0).

### 2.4. Animal sampling

In experiments 1 and 2, blood samples were collected from the infected animals at −1, 3- and 5-days post-inoculation (dpi) and from the sentinel animals at 0, 4-, 7-, 10- and 14-days post exposure (dpe) (Experiment 1, Figure 1). Nasal swabs were also collected during experiment 2 from the infected animals on −2, 3 and 5 dpi and from the sentinels at 9 and 14 dpe and placed in 1 ml DMEM with 2% FCS. All samples were stored at −80°C until analysis.

In experiment 3, unstabilized blood (to obtain serum), EDTA-stabilized blood (EDTA blood) and oral-, nasal-, and rectal swabs were collected prior to inoculation at 0 dpi and at 3, 5 and 6 dpi (euthanasia) (Figure 2). Urine samples were obtained on an occasional basis, i.e., if the pigs urinated while the personnel were in the pens, urine was collected with a tube during urination. Prior to their introduction into the three contaminated pens in boxes 4-6, unstablized blood, EDTA blood and oral-, nasal-, and rectal swabs were collected from the sentinel pigs. After introduction into the contaminated pens, blood and swabs samples were collected from the twelve pigs as shown in Figure 2.

### 2.5. Environmental sampling

#### 2.5.1. Pilot studies to test detection of ASFV DNA on spiked surfaces

The efficacy of recovering virus from a smeared surface and from straw was tested in pilot experiments. Samples (50 µl) of ASFV (strain BA71V) with titres of 10^4^, 10^5^, 10^6^ TCID_50_/ml were spiked directly onto the surface of petri dishes and smeared over the surface or dropped onto small clumps of straw of approx. 1 cm^3^. The surfaces were briefly allowed to dry and then swabbed either using pieces of normal household electrostatic dust swabs (Minky Homecare, Rochdale, UK) with an approximate size of 4 × 4 cm, or round tip traditional cotton swabs. Both types of swabs were placed into 1.5 ml of either culture medium (DMEM with 10% FBS and Penicillin/streptomycin) or PBS and kept for 2h at 4⁰C to elute the virus. The spiked straw clumps were also placed directly into the same volumes of either PBS or culture medium. Nucleic acids were extracted from 140 µl of each sample and also from the original virus dilutions with the “QIamp viral RNA extraction kit” (Qiagen) and eluted in 60 µl elution buffer according to manufacturer’s instructions. Viral DNA was detected by PCR from 2 µl of each sample using primers for the ASFV B646L (VP72) gene (CTGCTCATGGTATCAATCTTATCGA, GATACCACAAGATCRGCCGT, 200 nM), Platinum Blue PCR SuperMix (Invitrogen) in total volumes of 20 µl. The PCR program: 3 min. 94⁰C; 35 cycles of 20 sec 94⁰C, 20 sec 58⁰C, 20 sec 72⁰C was used. Amplification products were visualized using 1.5% agarose gel electrophoresis.

#### 2.5.2. Sampling of animal premises

In premises with infected animals in experiments 1 and 2, dust swabs were used to sample different surfaces, i.e., walls, floor, bedding (especially showing evident presence of animal excretions) and the rim of water bowls. The blood collection points of some of the infected animals were also sampled as positive controls of the swabbing technique and recovery of viral DNA. In animal experiment 1, dust swab areas were approximately 50 cm^2^ (5 × 10 cm) and after collection the samples were kept refrigerated at 4⁰C until transport to the lab and then immersed in 4 ml PBS at 4⁰C for 2h. DNA was extracted as described above using the “QIamp viral RNA extraction kit” (Qiagen) in duplicate from each swab. In animal experiment 2, a similar sampling method was followed but dust swab sizes were reduced to approximately 20 cm^2^ (4 × 5 cm) and immersed in 2 ml PBS at 4⁰C for 2h. DNA was extracted from 100 µl of each sample in duplicate using the automated extraction “Kingfisher Flex Extraction System” and Magvet Universal Isolation kit (Thermo Fisher Scientific, LSI MV384).

In the pens housing the infected pigs in experiment 3, floor swabs and faeces were collected from 0 dpi (before inoculation) and until the day of euthanasia (6 dpi). In addition, these samples were obtained on the day of introduction of the sentinel pigs (0 dpe) in each of the three pens. Floor swab samples were collected in 1 ml 1x phosphate buffered saline (1x PBS) (Thermo Fisher Scientific). Faecal homogenate suspensions (10%, w/v) were prepared in 1x PBS with 5% foetal bovine serum (FBS), streptomycin (Sigma-Aldrich), neomycin (Sigma-Aldrich), amphotericin (Sigma-Aldrich) and benzylpenicillin (Sigma-Aldrich). They were homogenized by rigorous vortexing with glass beads (Merck Millipore) and centrifuged at 950 x g for 10 minutes. Floor swabs in 1x PBS were vortexed and centrifuged briefly. Recovered supernatants were used for DNA extraction (see section 2.6).

#### 2.5.3. Air sampling of animal premises

Two systems were used to collect air samples during animal experiment 2. One was a handheld AirPort MD8 (Sartorius, Epsom, UK) that was used to collect samples from directly above the pigs for 5 min at a flow rate of 50 l min^−1^. Aerosol particles are retained on a gelatin filter (nominal pore size 3 μm) attached to the front of the device, through which air is drawn. The gelatin filter was dissolved in 10 ml RPMI with 10% FCS and penicillin and streptomycin after sampling. The second was a wet-walled cyclone Coriolis micro air sampler (Bertin Technologies, Aix-en-Provence, France) that was used to collect air samples from the room during husbandry and sampling. The Coriolis sampler was placed at a height of 1.1 m close to the extraction vent for the room, run for 30 minutes at a flow rate of 300 l/min and aerosolised material was collected into 10 ml of RPMI medium with penicillin and streptomycin and 10% FCS. Air samples were stored at −80⁰C until nucleic acids extraction from 100 µl of each, in duplicate, using the automated extraction “Kingfisher Flex Extraction System” and Magvet Universal Isolation kit (Thermo Fisher Scientific, LSI MV384).

### 2.6. ASFV genome detection by qPCR

For animal experiments 1 and 2, nucleic acids extracted from environmental samples as described above (2.5.2) were analysed for the presence of ASFV DNA by qPCR as described previously (King, Reid et al. 2003). Extracted nucleic acids (5 µl) were tested per sample in duplicate. ASFV DNA quantification in whole EDTA blood collected from the animals was performed similarly after extraction in duplicate from each blood sample using the extraction system Magvet Universal Isolation kit (Thermo Fisher Scientific, LSI MV384) and automated extraction with a Kingfisher Flex Extraction System as described above.

In experiment 3, DNA was purified from EDTA blood, nasal-, oral-, -rectal- and floor swab samples, urine, and faecal supernatants using a MagNA Pure 96 system (Roche) and analysed for the presence of ASFV DNA by qPCR (essentially as described in (Tignon, Gallardo et al. 2011) and (Olesen, Lohse et al. 2017) but using the Bio-Rad CFX Opus Real-Time PCR System (as previously, Olesen, Lazov et al. 2023). Results are presented as viral genome copy numbers (per mL: EDTA blood, swab supernatant, urine, or per gram: faecal suspension supernatants) using a standard curve based on a 10-fold dilution series of the pVP72 plasmid (Olesen, Kodama et al. 2021). A positive result in the qPCR was determined to be a threshold cycle value (Cq) at which FAM (6-carboxy fluorescein) dye emission increased above background within 42 cycles (as previously, Olesen, Lazov et al. 2023.

### 2.7. Detection of infectious ASFV using virus isolation in cells

Aliquots of nasal swabs and of air samples collected during animal experiment 2 were inoculated onto primary porcine bone marrow cell cultures. A sample (0.8 ml) of each nasal swab (out of 1 ml total) and 1 ml of each air filter sample (out of 10 ml total volume) were added to the cells cultivated in 6-well plates with RPMI medium containing 10% FBS and penicillin/streptomycin in 3 ml culture volume. The cells were then incubated and observed for development of hemadsorption for a period of 5 days after which the plates were frozen at −80°C. After thawing and centrifuging at 600 ×g for 5 minutes, aliquots of 1 ml of supernatant from each of the first inoculation wells were used to inoculate new primary cultures. These were again incubated and observed for 5 days for development of hemadsorption.

In experiment 3, swab samples, urine and faecal suspension supernatants were analysed for the presence of infectious ASFV by virus isolation in PPAM (Olesen, Lohse et al. 2017, Olesen, Lohse et al. 2018). The cells were maintained in Minimum Essential Medium (MEM, Gibco) with 5% FBS in NUNC 24-well plates (Thermo Fisher Scientific). Prior to inoculation of cells, PBS from the swab samples, faecal suspensions and urine samples were filtered, using0.45 µm syringe filters (Merck Millipore) and the clarified samples (100 µl) were added to MEM (100 µL) containing antibiotics and 10% FBS prior to addition to 1 mL PPAM (2 x 10^6^ cells/mL). In one trial, the inoculum was removed from the cells after incubation at 37°C for 1h, and the cells were then washed twice with PBS. MEM, containing 5% FBS, streptomycin, neomycin, amphotericin and benzylpenicillin, was added to the cells and incubated at 37°C (5% CO_2_) for three days. In another trial, PPAM (2 x 10^6^ cells/mL with 5% FBS) were incubated with the inoculum for three days, i.e. without its removal, at 37°C (5% CO_2_). In both trials, following the three days, the cells were harvested by freezing and 100 µL of the harvested 1^st^ passage was inoculated onto 1 mL fresh PPAM (2 x 10^6^ cells/mL with 5% FBS) in NUNC 24-well plates. Following three days of incubation (37°C, 5% CO_2_), virus-infected cells were identified using an immunoperoxidase monolayer assay (IPMA) essentially as described previously (Botner, Nielsen et al. 1994, Olesen, Lohse et al. 2017). Red-stained (virus-infected) cells were identified under a light microscope.

### 2.8. Antibody detection

Blood from the sentinel animals at termination of experiment 2 was tested for the presence of anti-ASFV antibodies using lateral flow test devices (INGEZIM PPA CROM, R.11.PPA.K41, Ingenasa). Serum samples obtained at euthanasia from the inoculated and the sentinel pigs in experiment 3 were tested for the presence of anti-ASFV antibodies using the INgezim PPA Compac kit (Ingenasa) according to the manufacturer’s instructions.

## 3. Results

### 3.1. Animal experiments and result of naïve pig exposures

The aim of the experiments was to assess the risk that environmental contamination poses for infection of naïve pigs introduced to contaminated pens at different days after infected pigs were removed. In experiment 1, a group of five pigs (numbered 41 to 45) were inoculated with 10,000 HAD_50_ infectious units of virulent ASFV isolate by the intramuscular route (0 dpi). Clinical signs typical of acute ASF including increased temperature rising above 41°C, anorexia and increasing lethargy were detected from 3 dpi (Fig 3a, b) but no clear haemorrhagic faeces or other haemorrhagic lesions or excretions were observed. All pigs reached the predefined humane endpoint at 5 dpi and were culled. Measurement of viremia by qPCR confirmed the expected high levels (above 10^8^ genome copies per ml of blood) by 5 dpi (Fig. 3c). The room was then left completely uncleaned for one day and two sentinel pigs (numbered 46, 47) were introduced into the room. Over the period of 14 days, until the end of the experiment, no clinical signs of ASF were observed in the sentinel pigs (Fig. 3d, e). Blood samples collected along this period also showed no detectable ASFV genomes in either of the animals. Thus, neither of the sentinels became infected and environmental transmission did not occur.

**Figure 3:**
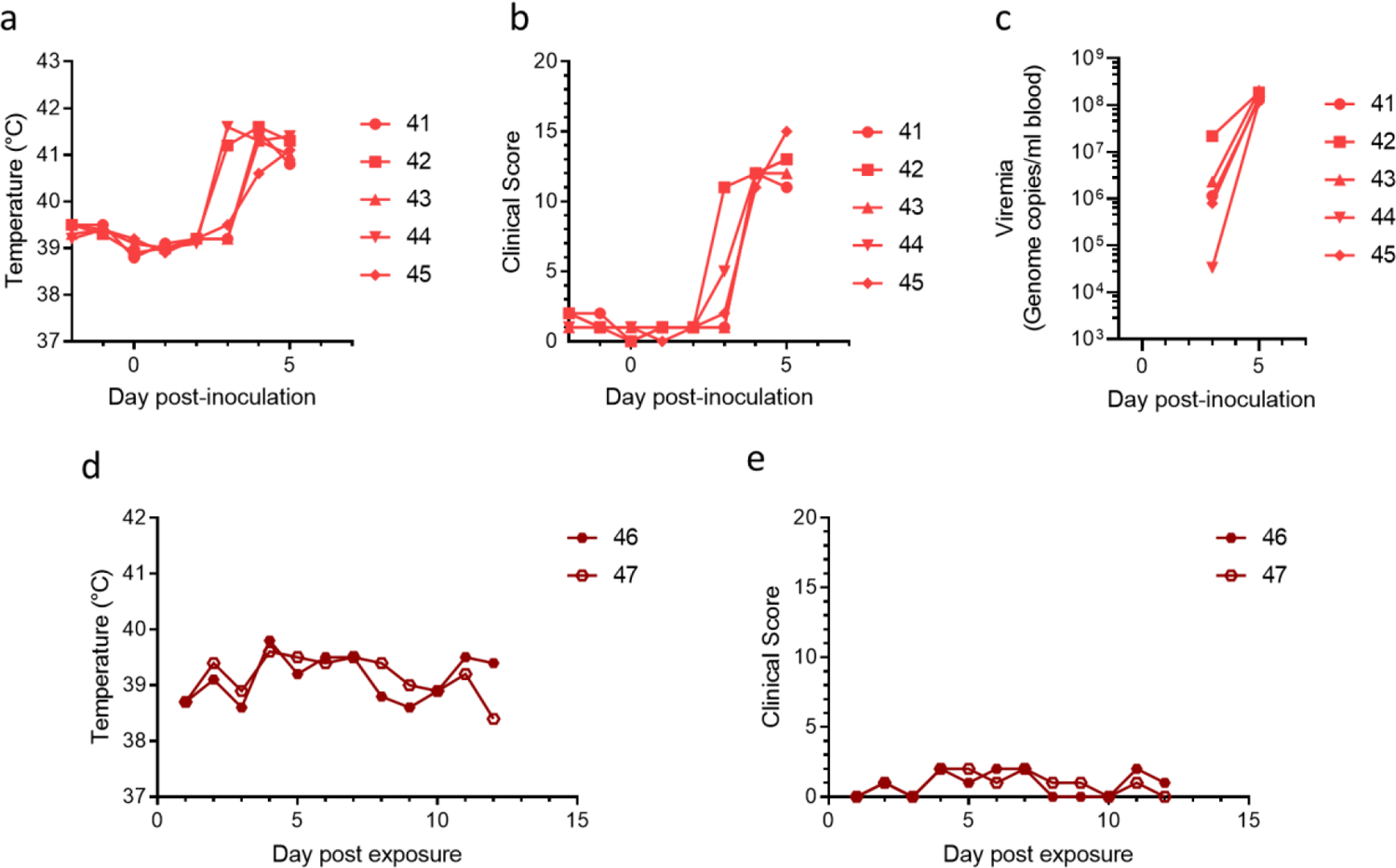
Clinical parameters for inoculated animals (a-b) and sentinels (d-e) and viremias of the inoculated animals (panel c) during Experiment 1.

In experiment 2, we shortened the time between removal of infected animals and introduction of sentinels. Two sentinel animals were introduced on the same day as a group of six infected animals with virulent ASFV were removed due to reaching the predefined humane endpoint. All six of the directly inoculated pigs developed clinical signs from 3 dpi and were culled at 5 dpi, as in experiment 1 (results not shown). All animals had very high blood viremias as detected by qPCR on day 5 with titres ranging from 10^7.69^ - 10^9.11^ (Table 1). We could clearly detect viral DNA in nasal swabs at this timepoint, with 10^5.39^ to 10^6.16^ copies per swab, but not two days earlier, i.e. on day 3. Infectious virus was isolated from the nasal swabs of two animals, one at day 3 and another at day 5 post-infection (Table 1 and Figure S2 in supplementary material). However, in experiment 2 again no clinical signs or viremia were detected in the sentinel pigs during the 15-day period of exposure, and no ASFV specific antibodies were detected in the blood of the animals either. Nasal swabs collected at 9 dpe and 14 dpe were also negative for viral DNA (not shown). Thus, in experiment 2 transmission from environmental contamination to naïve pigs also failed when these were exposed to the contaminated environment on the same day acutely infected animals were removed from the premises.

**Table 1:** ASFV genome copy numbers in blood and in nasal swabs of inoculated animals during animal experiment 2 and presence of infectious virus in nasal swabs. Genome copy numbers were determined by qPCR and represent the average for each sample tested in duplicate. Virus isolation was performed in primary macrophage cell cultures. “n.d.” - not detected; “n.a.”-not available (nasal swab was not collected).

|  | 3 dpi |  |  | 5 dpi |  |  |
| --- | --- | --- | --- | --- | --- | --- |
|  | Genome<br>per ml of<br>blood<br>(Log<sub>10</sub>) | Genome<br>per nasal<br>swab<br>(Log<sub>10</sub>) | Infectious<br>virus in<br>nasal<br>swabs | Genome<br>per ml of<br>blood<br>(Log<sub>10</sub>) | Genome<br>per nasal<br>swab<br>(Log<sub>10</sub>) | Infectious<br>virus in<br>nasal<br>swabs |
| **Pig 1** | 7.79 | n.d. | no | 9.11 | 6.13 | no |
| **Pig 2** | 6.55 | n.d. | no | 8.80 | 6.16 | yes |
| **Pig 3** | 7.99 | n.d. | yes | 8.75 | n.a. | n.a. |
| **Pig 4** | 4.61 | n.d. | no | 8.37 | 5.70 | no |
| **Pig 5** | 5.45 | n.d. | no | 8.39 | 5.86 | no |
| **Pig 6** | 4.09 | n.d. | no | 7.69 | 5.39 | no |

In experiment 3, pigs 1–12 in boxes 4-6 (see Figure 2) were inoculated intranasally with a virus suspension containing 10,000 TCID_50_/2 ml. At 4 dpi, three out of four inoculated pigs in boxes 4 (pigs 1, 2 and 4) and 6 (pigs 10, 11 and 12) presented with high fever (rectal temperature above 41 °C). In box 5, two out of four inoculated pigs presented with high fever at 5 dpi (pigs 7 and 8). Clinical signs became apparent from 4 dpi (boxes 4 and 6) or 5 dpi (box 5) and included depression, anorexia, mildly laboured breathing, hyperemia of the skin and cyanosis on the ears and distal limbs, blood in faeces (pig 10, box 6 at 6 dpi, one day before introduction of sentinel pigs into this pen) and vomiting (pig 8, box 5). At 6 dpi, pig 2 was found dead upon entering box 4 (so no clinical score was registered on this day for this pig). Foam was observed from the nostrils of this pig. The remaining 11 inoculated pigs were euthanized on this day. Pigs 1, 4 (box 4), 7, 8 (box 5), 10, 11 and 12 (box 6) had reached the pre-determined humane endpoints. The remaining four, pigs 3 (box 4), 5, 6 (box 5) and 9 (box 6) were euthanized for animal welfare reasons. Rectal temperatures and clinical scores for the inoculated pigs are shown in Figure 4a, b. All but pigs 3 (box 4) and 5 (box 5) had shown clinical signs of ASF. Measurement of viremia by qPCR confirmed the expected high levels (above 10^8.7^ genome copies per ml of blood) by 6 dpi in all animals except pig 3 in box 4 that had approximately 10^5^/ml and pig 5 in box 5 that did not show viremia (Figure 4c). Levels of viral DNA detected in nasal, oral, and rectal swabs obtained from the inoculated pigs at 6 dpi are shown in Table 2 A-C. Most pigs in box 4 (pigs 1, 2, 4) had high levels of viral genome in nasal swabs with at least 10^8^ copies/ml (pig 3 had 10^4.4^), oral swabs with 10^4.6-7^ and rectal swabs with 10^6.2-7.2^ (except again for pig 3). Pigs in box 5 (pigs 5-8) had lower ASFV DNA levels in nasal swabs than the previous group, with no viral DNA in one of the animals and 10^4.8-7.8^ copies/ml in the other three; only one animal had viral DNA in its oral swab with 10^5.1^ and three of the animals had rectal swabs with 10^3.7-6.3^ copies/g. All pigs in box 6 (pigs 9-12) had quite high levels of ASFV DNA in nasal swabs, with 10^6.8-8.6^ copies/ml, oral swabs with 10^6.2-7.1^/ml and rectal swabs with 106.2-7.5/ml.

**Figure 4:**
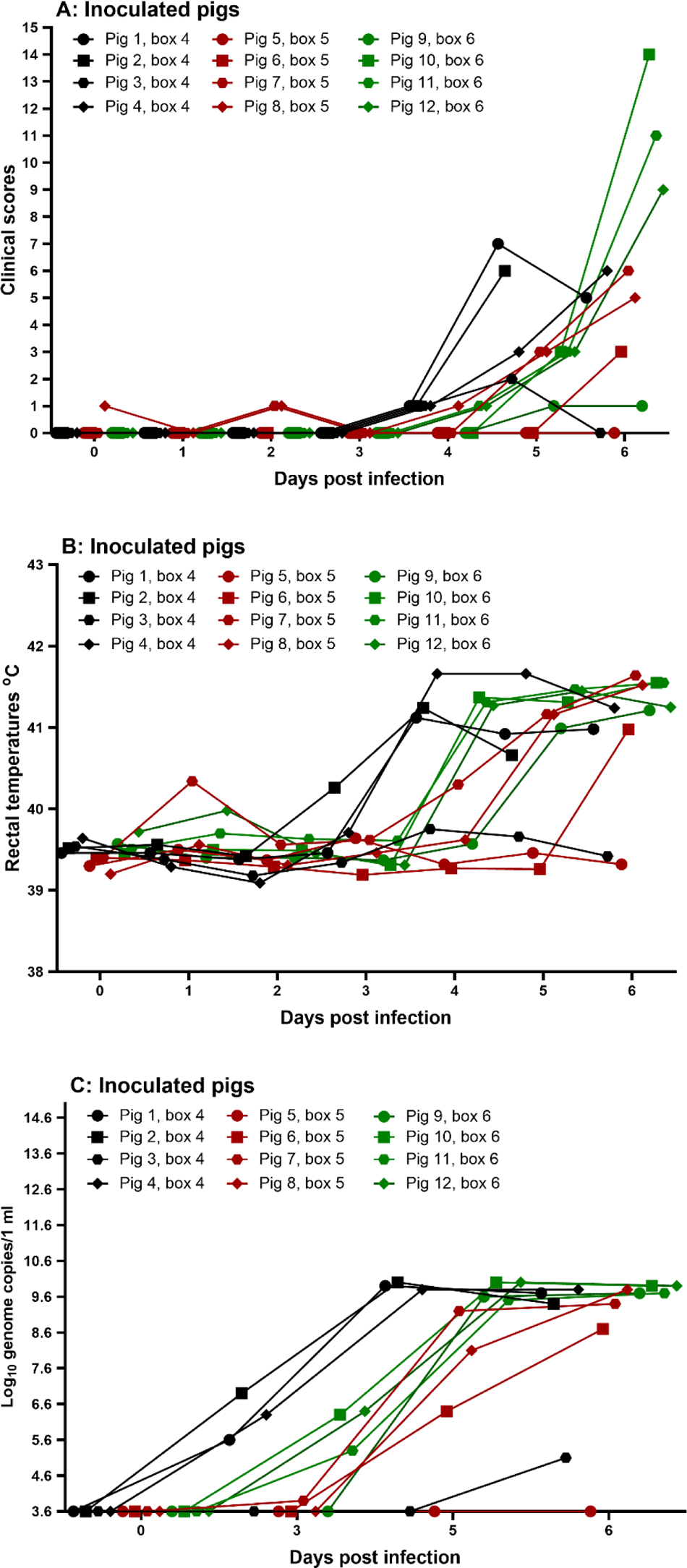
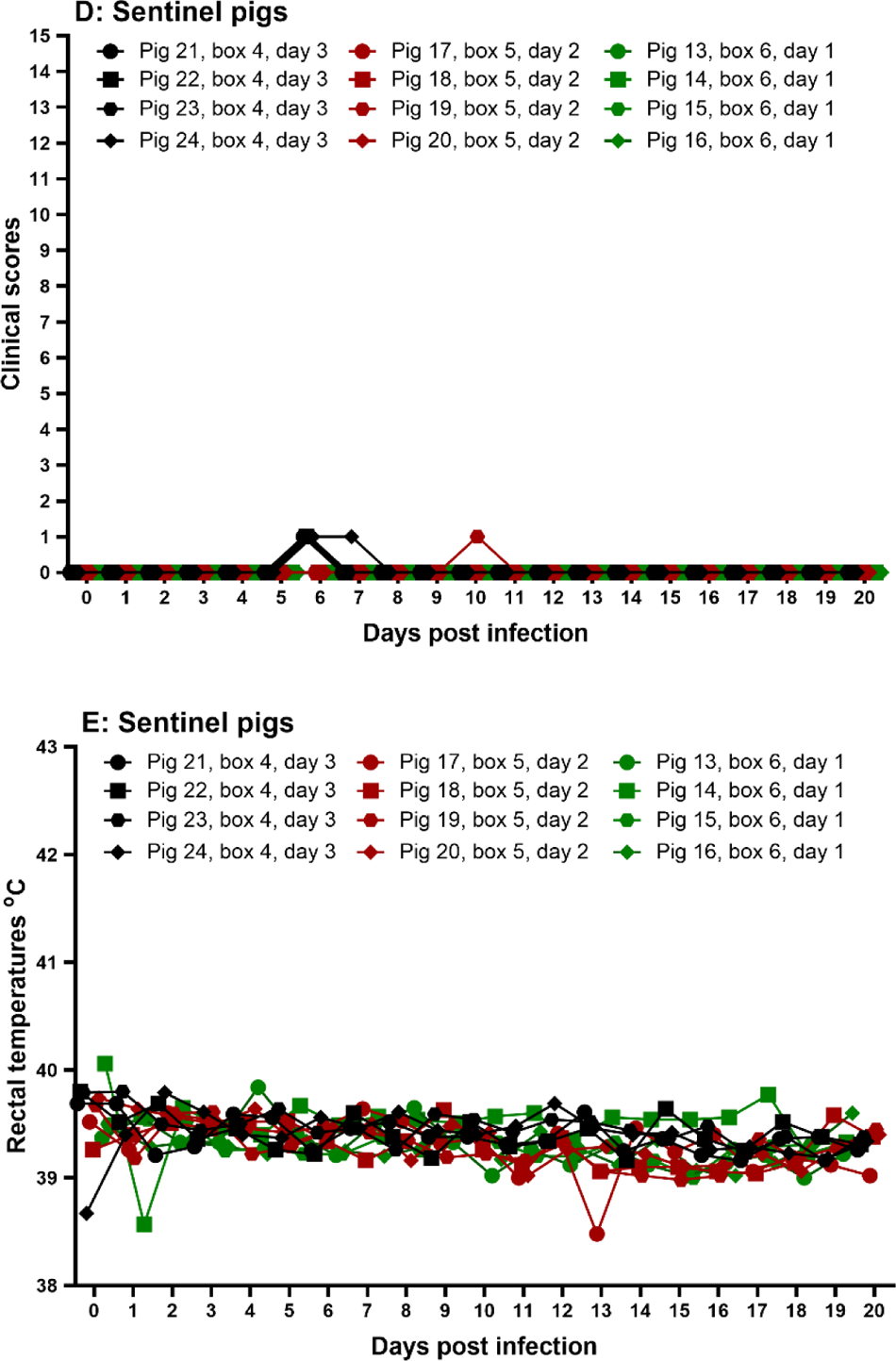
Clinical parameters from the inoculated and sentinel pigs in Experiment 3. Inoculated pigs’ clinical scores (A), rectal temperatures (B), detection of ASFV DNA in EDTA-blood (C) and sentinel’s clinical scores (D) and rectal temperatures (E). In panel C, the detection threshold for ASFV DNA is 10^3.6^ genome copies/ml. The data on clinical scores and rectal temperatures from pigs 9-12 has been published previously for a different study (Olesen, Kodama et al. 2021) and is shown here for c<u>ompleteness</u>.

**Table 2:**
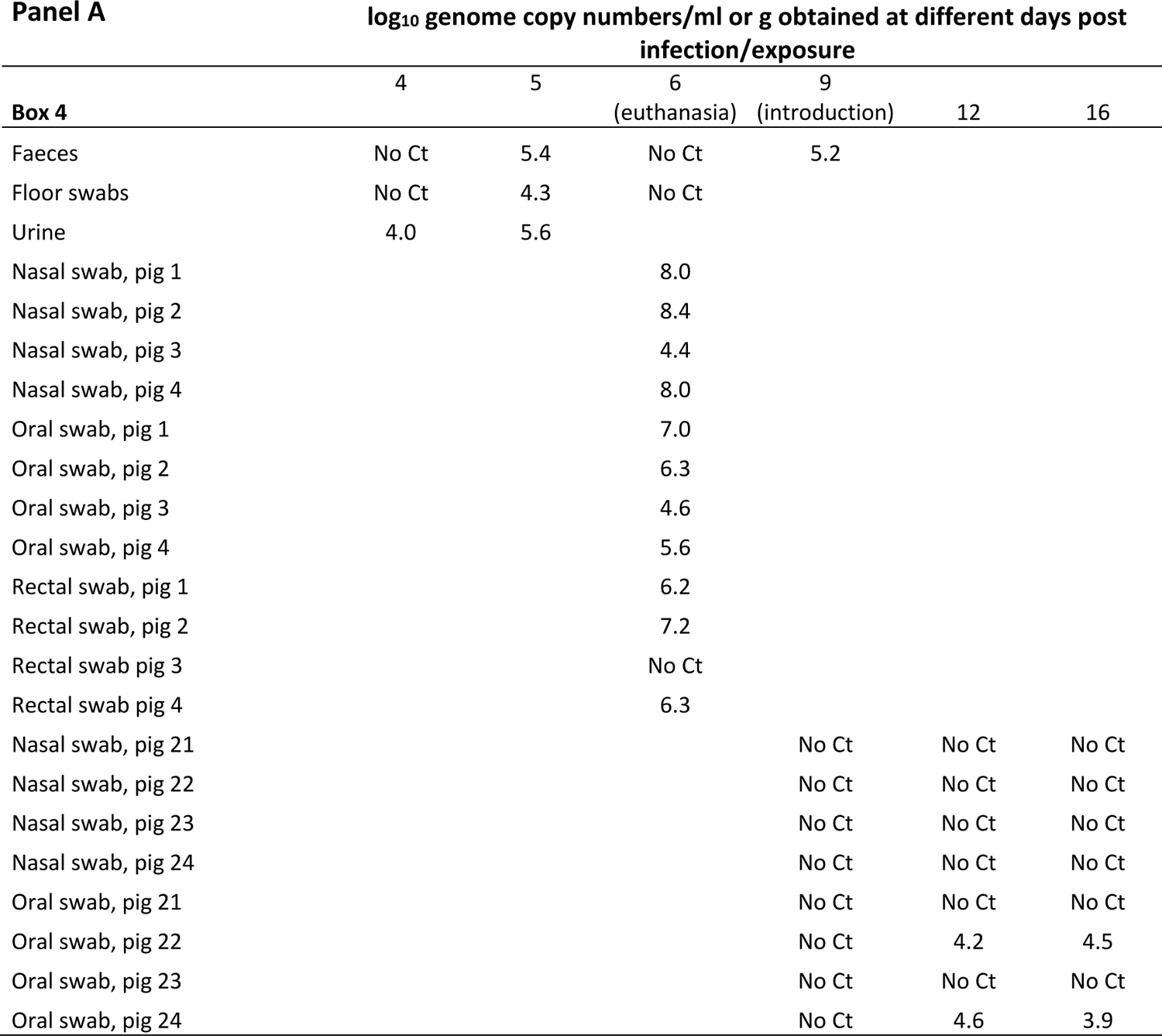
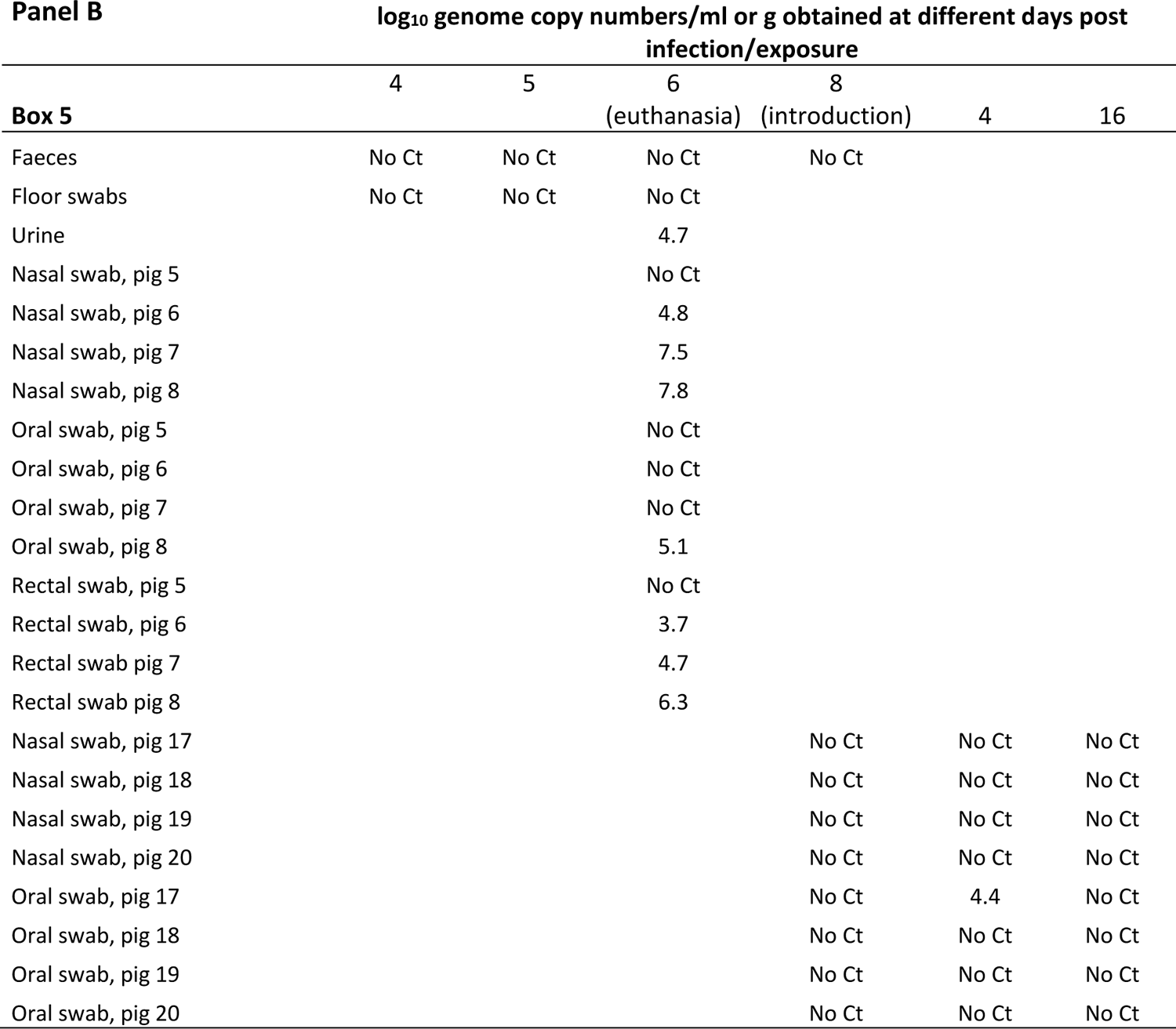
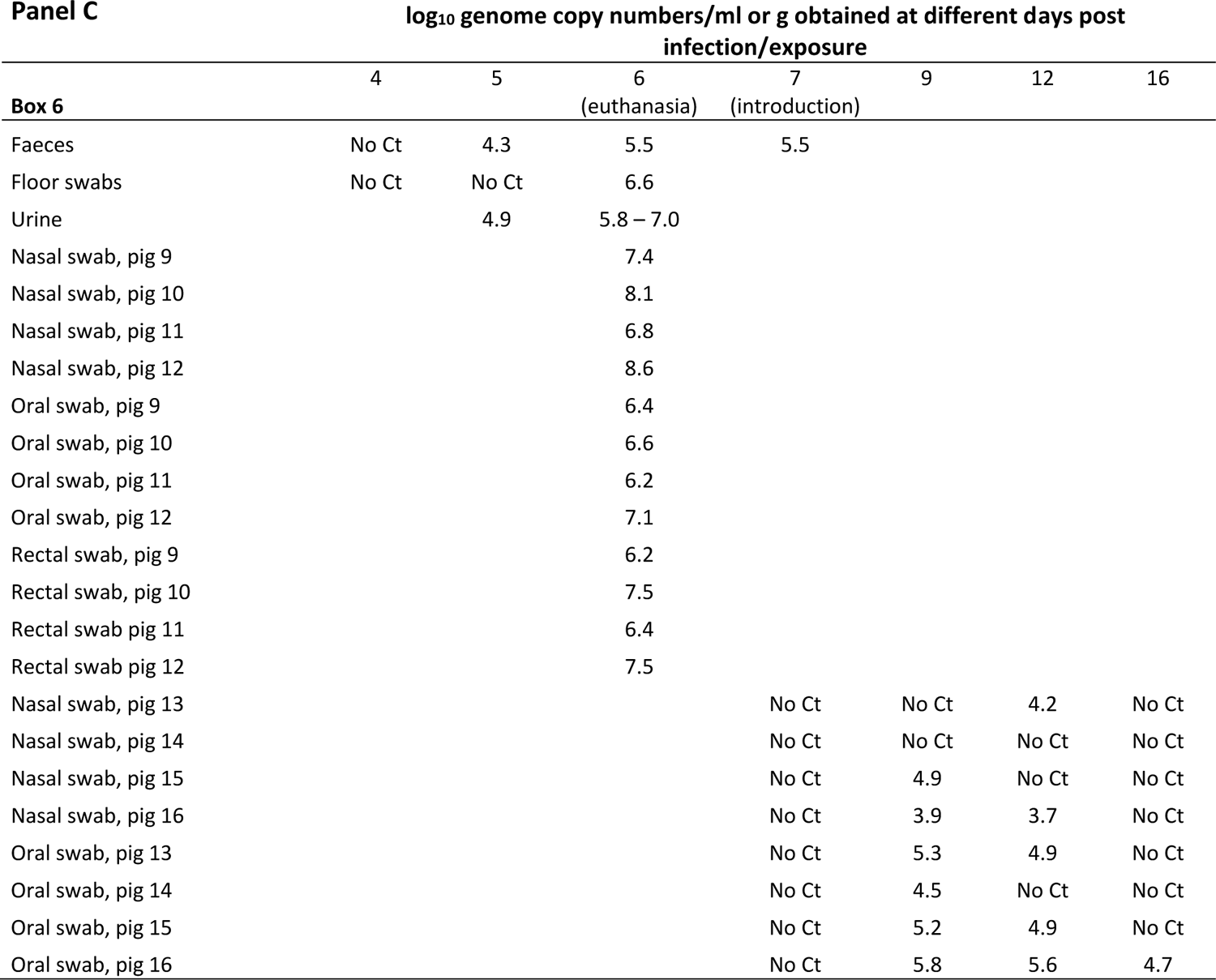
qPCR results for ASFV DNA detection in faecal and urine samples and in floor, nasal, oral and rectal swabs obtained in the different boxes during Experiment 3. Sentinel pigs were introduced to the contaminated environment at different days following euthanasia of the infected pigs: 3 days in box 4 (panel A), 2 days in box 5 (panel B) and 1 day in box 6 (panel C). Swab samples at 0 dpe were collected prior to the introduction of the pigs into the contaminated environment in the three boxes. Numbers are log_10_ genome copy numbers/ml (swab samples and urine) or log_10_ genome copy numbers/g (faeces).

Following exposure to the contaminated environments after 1 day (box 6), 2 days (box 5) or 3 days (box 4) following removal of the infected pigs, none of the sentinels (pigs 13-24, four per box), developed clinical signs that would indicate an ASFV infection. Rectal temperatures and clinical scores obtained from these pigs are shown in Figure 4d-e.

No ASFV DNA was detected in EDTA-blood obtained from the sentinel pigs (data not shown). After their introduction to the contaminated environment, ASFV DNA was detected in several nose and mouth swabs (Table 2 A-C). The highest prevalence and most viral DNA in swabs were observed in the sentinel pigs that were introduced into box 6 one day after euthanasia of inoculated animals. At 2 dpe all oral swabs from pigs in this group were positive for ASFV DNA (10^4.5-5.8^ genome copies) and viral DNA was also detected in the nasal swabs (10^3.9-4.9^) from two of the animals. At 5 dpe however, there was a reduction in the number of positive swabs and by 9 dpe only one of the animals had a positive oral swab with 10^4.7^ genome copies (Table 2 C). Following introduction to box 5, at day 2 after the infected animals were removed, only one sentinel had a positive oral swab at 4 dpe with 10^4.4^ genome copies and no swabs were positive at 8 dpe (Table 2 B). In box 4, where sentinels were introduced at day 3, ASFV DNA was only detected in oral swabs of two of the animals at both 3 and 7 dpe with 10^3.9-4.6^ genome copies (Table 2A). Despite the apparent uptake of ASFV by the sentinel animals from the contaminated environment, as evidenced by the presence of viral DNA in nasal and oral swabs at least at early days post exposure, all sentinel pigs were euthanized after three weeks exposure to the contaminated environment without evidence of infection by ASFV. No ASFV specific antibodies were detected in the blood of the animals after the 21 days exposure.

### 3.2. Estimation of levels of virus contamination in environmental samples in rooms housing infected pigs

Pilot experiments were carried out to compare swabbing methods to recover virus from surfaces. In these experiments 50 µl of ASFV containing 5 × 10^4^, 5 × 10^3^ and 5 × 10^2^ TCID_50_ was spiked directly onto petri dishes or onto small clumps of straw. The surfaces were then swabbed and ASFV DNA was detected by PCR. PCR fragments could be weakly detected only from spiking with 5×10^4^ TCID_50_ collected with either dust or cotton swabs or directly from straw. PBS was the most effective for elution. Control samples of ASFV DNA gave clear positive results (See Figure S1 in Supplementary material). This showed that the sample swabbing method using dust swabs and PBS for viral elution was sufficiently sensitive to detect 5×10^4^ TCID_50_ of virus spiked on straw and on surfaces by conventional PCR.

The premises housing ASFV infected animals are typically contaminated via the different animal excretions and sometimes blood. Animals experimentally infected (by the intramuscular route) with high virulence isolates of ASFV develop clinical signs of acute disease between day 3 and 5 after infection concomitant with excretion of infectious virus (Guinat, Reis et al. 2014) (Davies, Goatley et al. 2017). Previously we detected infectious virus and viral DNA in urine and faeces collected from infected animals at the onset of pyrexia (≥ 40°C) (Davies, Goatley et al. 2017). In urine, we could detect infectious virus at approximately 10^3^ TCID_50_/ml and viral genome at 2.5 × 10^4^ copies/ml. In faeces, virus was intermittently detected but was present in some samples at up to 6.8 × 10^4^ TCID_50_/g or 10^7^ genomes/g. In rectal swabs taken from animals after the onset of clinical signs, between days 3-6 after infection, up to 10^2^ HAD_50_/ml and 10^3-4^ genome copies/ml could be detected and in nasal swabs, up to 10^4^ HAD_50_/ml and 10^5^ genome copies/ml (Guinat, Reis et al. 2014). Infectious virus was not detected in oral swabs although low levels of genome could be detected. In blood, much higher levels of virus and genome were detected, up to 10^6-8^ TCID_50_/ml and 10^6-8^ genome copies/ml (Guinat, Reis et al. 2014, Davies, Goatley et al. 2017). Therefore, the swabbing technique and qPCR detection of ASFV genome should be sufficiently sensitive to detect contamination with these secretions and excretions and especially with blood, if present in the areas swabbed.

During animal experiments 1 and 2, we assessed the recovery of ASFV DNA from roughly 20 cm^2^ areas sampled with dust swabs in the premises housing ASFV infected animals. A similar sampling regime was followed as used during previous experiments with foot-and-mouth disease virus (FMDV) in which FMDV nucleic acid (RNA) was readily detected in most samples (Brown, Nelson et al. 2021). During animal experiment 1, the room housed five ASFV infected pigs. Two swabs each were collected from the floor, wall, straw bedding, and water bowl at day 1 before infection and days 3 and 5 post-infection with virulent ASFV OURT88/1, on the same days that blood samples were collected from each of the inoculated animals. The animals were euthanised on day 5 at the humane end point. As control for the recovery of ASFV DNA by the swabbing technique, the skin area surrounding the site of blood collection from two of the animals at days 3 and 5 post-infection was also swabbed. Swabs were also collected from the environment following removal of the ASFV infected pigs and introduction of the sentinel pigs, on days 6 and 7 post-inoculation (or 0 and 1 dpe). Using the swabbing method, a low amount of ASFV DNA (500 - 750 genome copies) was detected by qPCR in eluates from the swabs of a few of the environmental surfaces at days 5 and 6 post-infection: on day 5 in one swab from the floor and one from a wall, and on day 6 in both swabs from straw bedding (Table 3, Experiment 1). None of the swabs from day 7 or from the day before infection showed detectable ASFV DNA (not shown). ASFV DNA was always detected in positive control samples (swabs around the site of blood collection from pigs): approximately 10^4^ and 10^5^ genomes on day 3 rising to 4 and 8 × 10^7^ on day 5. The genome copy numbers per ml determined directly from the blood of the infected animals by qPCR ranged between 10^4^-10^7^ at day 3 and was approximately 10^8^ at day 5 (Table 3, Experiment 1 and Fig. 2c). The difference was less than 1 log_10_ in genome copies detected between the swabs from areas around the sites of pig bleeding and the blood samples on day 5 but this is likely to be due to lower recovery of DNA from the swabs. Detecting ASFV DNA in environmental samples with lower viral loads may be difficult and require a larger sampling area.

**Table 3.**
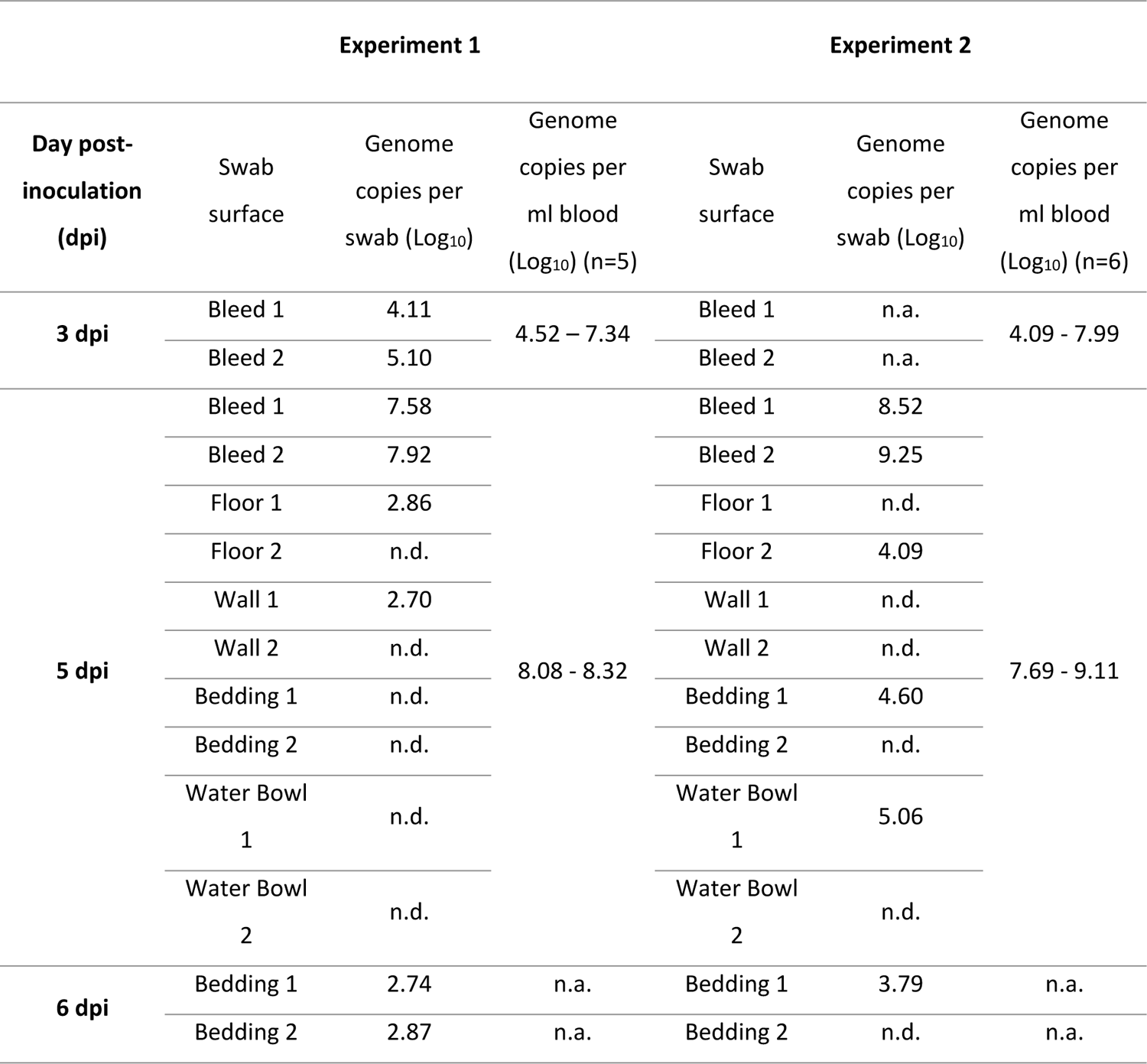
ASFV genome copy numbers in the environment of premises with infected animals and in the blood of the animals during Experiments 1 and 2. Swabs were collected from different surfaces of the premises or from areas around bleeding of two of the infected pigs (Bleed 1 and 2). Total genome copy numbers of each swab were determined by qPCR and represent the average for each sample tested in duplicate. For comparison, the levels ASFV genome in the blood of the animals occupying the premises are also shown. Genome copies per ml of blood represent the range of values determined in the infected group of pigs. “n.a” not available, “n.d.” not detected.

| Experiment 1 |  |  |  | Experiment 2 |  |  |
| --- | --- | --- | --- | --- | --- | --- |
| Day post-inoculation (dpi) | Swab surface | Genome copies per swab (Log<sub>10</sub>) | Genome copies per ml blood (Log<sub>10</sub>) (n=5) | Swab surface | Genome copies per swab (Log<sub>10</sub>) | Genome copies per ml blood (Log<sub>10</sub>) (n=6) |
| 3 dpi | Bleed 1 | 4.11 | 4.52 – 7.34 | Bleed 1 | n.a. | 4.09 - 7.99 |
|  | Bleed 2 | 5.10 |  | Bleed 2 | n.a. |  |
| 5 dpi | Bleed 1 | 7.58 | 8.08 - 8.32 | Bleed 1 | 8.52 | 7.69 - 9.11 |
|  | Bleed 2 | 7.92 |  | Bleed 2 | 9.25 |  |
|  | Floor 1 | 2.86 |  | Floor 1 | n.d. |  |
|  | Floor 2 | n.d. |  | Floor 2 | 4.09 |  |
|  | Wall 1 | 2.70 |  | Wall 1 | n.d. |  |
|  | Wall 2 | n.d. |  | Wall 2 | n.d. |  |
|  | Bedding 1 | n.d. |  | Bedding 1 | 4.60 |  |
|  | Bedding 2 | n.d. |  | Bedding 2 | n.d. |  |
|  | Water Bowl 1 | n.d. |  | Water Bowl 1 | 5.06 |  |
|  | Water Bowl 2 | n.d. |  | Water Bowl 2 | n.d. |  |
| 6 dpi | Bedding 1 | 2.74 | n.a. | Bedding 1 | 3.79 | n.a. |
|  | Bedding 2 | 2.87 | n.a. | Bedding 2 | n.d. | n.a. |

In experiment 2, swab sizes and elution volumes were reduced to approximately half to concentrate the potential ASFV DNA from contaminated surfaces and increase sensitivity of detection. Swabs were collected from the premise’s surfaces on days 3, 5 (0 dpe) and 6 (1 dpe), and on day 5 also from the blood collection points on the animals. This time, viral DNA levels detected on the premise’s surfaces were higher than in experiment 1, in the range of 10^3.8^ to 10^5^ copies on days 5 and 6, detected on the floor, bedding and a water bowl (Table 3, Experiment 2). Increased genome copy levels were also present in the blood of some of the infected animals in comparison to experiment 1 at day 5 dpi, which may have led to more virus being released to the environment. The more concentrated elution of the swabs may also have contributed to the higher levels of detection of ASFV DNA during experiment 2. Air samples were also collected from the premises during the second experiment, on 0, 3, 5 and 6 dpi using two different devices (MD8 and Coriolis). No viral DNA or infectious virus was detected on any of the sampled days, suggesting that ASFV was not aerosolized to a high level during the experiment and may have been efficiently removed by the ventilation system.

In experiment 3, viral DNA was also detected in urine and faeces excreted into the pens housing the infected pigs and in floor swabs (see Table 2 A-C). The level of contamination of the three boxes seemed to differ. Viral DNA was detected in floor swabs and in urine and faecal samples obtained prior to or on the day of introduction of sentinel pigs into boxes 4 (3-day group) and 6 (1-day group), but in box 5 (2-day group) only one urine sample was positive 2 days before introduction. The swab samples from the inoculated animals in box 5 also showed lower prevalence and levels of ASFV DNA than in the other two boxes and one of the animals was not viremic (see Fig. 4), which may explain the lower environmental contamination detected in this box. The level of ASFV DNA in the faecal samples and floor swabs in boxes 4 and 6 varied from 10^4.3^-10^5.5^ genome copies/g and 10^4.3^-10^6.6^ genome copies/mL, respectively. The level in urine obtained from the inoculated pigs in the three boxes varied from 10^4^-10^7^ genome copies/ml (Table 2 A-C). However, no infectious virus was detected in the urine samples, faeces supernatants or floor swabs following two passages in PPAM (not shown).

## 4. Discussion

In three separate transmission experiments we failed to detect transmission of ASF to naïve pigs introduced into rooms which recently had housed pigs with acute ASFV infection. From previous studies we knew that the amount of virus in urine, faeces and oral and nasal secretions is much lower than in blood from infected animals. Infected blood may be present in the environment from excretions or scratches on pigs making transmission more likely. At later stages of infection, bloody diarrhoea may be observed but it was rarely observed in our experiments probably because in experimental settings, pigs are usually culled for humane reasons before reaching these late (terminal) stages of the infection.

Overall, in the current study, in experiments 1 and 2 we detected only low levels of ASFV genome in environmental samples from surfaces including walls, floor, water bowls and straw bedding (10^2.7^ to 10^5^ genomes in swabs) on days −1 and 0 before exposure of sentinels, introduced to the premises one day after or on the same day as removal of acutely infected animals. This suggests very low level of virus contamination in the environment consistent with the lack of transmission from the environment to sentinel pigs. In experiment 3, susceptible pigs were introduced to ASFV contaminated pens either one, two or three days after removal of ASFV infected animals (boxes 6, 5 and 4 respectively). Environmental sampling in the pens before or on the day of introduction showed viral DNA present in faeces and floor swabs only in boxes 4 and 6 (10^4.3^-10^5.5^ genome copies/g and 10^4.3^-10^6.6^ genome copies/ml respectively), but not in box 5. ASFV DNA was detected in urine samples from all pens prior to introduction of sentinel pigs, with 10^4^-10^7^/ml genome copies. The level of contamination within the three pens seemed to depend on the course of infection in the nasally inoculated pigs housed within them. Hence, in box 5, in which only half of the inoculated pigs had a severe course of infection, contamination levels of the pen seemed to be lower, when compared to boxes 4 and 6, where more inoculated pigs showed severe symptoms of the disease. In addition, the level of contamination of the pens also correlated with the degree of detection of ASFV DNA in nasal and oral swab samples obtained from the sentinel pigs, especially at earlier days of exposure, denoting uptake of virus from the environment. Thus, in box 5, which seemed to be only mildly contaminated, viral DNA was only detected in one oral swab from one sentinel pig out of four, whereas in box 4 oral swabs from two sentinels were positive and in box 6 oral swabs from all four sentinels and additionally three nasal swabs were positive. Further, in box 6, one of the infected pigs was observed to have blood in faeces (at euthanasia, one day prior to the introduction of sentinel pigs) which probably contributed to the increased environmental contamination in this box. Even with the highest contamination level and earliest introduction of the sentinels, no ASFV transmission occurred in box 6.

In an earlier study, transmission to pigs via an environment contaminated with ASFV was observed when a contaminated pen had been left empty for three days, but not for five days (Montgomery 1921). Other previous experiments (Olesen, Lohse et al. 2018) showed that pigs that were introduced into the contaminated environment at 1 day after removal of infected animals with acute disease, developed clinical disease within 1 week, and both ASFV DNA and infectious virus were detected in their blood. However, pigs introduced into the contaminated pens after 3, 5 or 7 days did not develop signs of ASF and no viral DNA was detected in blood samples within the following 3 weeks. The results suggested a relatively narrow window of time for transmission, but further repetitions were needed to confirm this. This short window could also be related to the limited half-life of infectious virus in the environment. The authors reported clinical signs including mild rectal bleeding in one of the directly infected pigs occupying the pen before exposure of the healthy pigs when environmental transmission was observed, as well as presence of viral DNA in faeces in the environment 1 day before exposure. Especially the contamination of the environment with blood from an acutely diseased animal, which typically contains high titres of infectious virus, was probably a main factor for transmission. This and other environmental conditions, including humidity and rate of air changes, may explain differences between the earlier experiments and the current study. For example, we observed that drying of ASFV samples resulted in a 10-fold loss of virus titre (Flannery et al., unpublished results). In the studies demonstrating transmission of ASFV to pigs via a contaminated pen environment, hay (Montgomery 1921) or large amounts of straw (Olesen, Lohse et al. 2018) were left within the contaminated pens. One study did report that matrices with a low moisture content (hay, straw, grain) can provide a suitable environment to ensure ASFV viability when compared to matrices with a higher moisture content (soil water, leaf litter) when stored cooled temperatures (Mazur-Panasiuk and Wozniakowski 2020). Infectious ASFV was detected for up to 56 days in spleen tissue from ASFV-infected pigs incubated with straw and hay, and for up to 28 days in spleen tissue incubated with grain, when the samples were kept at 4°C. At room temperature, a rapid decay of the virus within all matrices was observed and no infectious virus was detected after incubation for seven days in hay, straw, or grain (Mazur-Panasiuk and Wozniakowski 2020). Perhaps ASFV transmission via contaminated pens after one day (Olesen, Lohse et al. 2018) or three days (Montgomery 1921) was also observed as a result of a stabilizing effect of the bedding material on the virus. In experiments 1 and 2 in the current study we also detected viral DNA in straw used as bedding material on the days of introduction of susceptible animals, although the levels were not high (10^2.7^ to 10^4.6^ genome copies). The environment into which the pigs were introduced in experiment 3 had no straw and had become very dry when compared to the environment that the pigs were introduced to in Olesen et al. (2018) (Olesen, Lohse et al. 2018), where a thick layer of straw prevented the environment from drying out. Furthermore, pigs introduced into an environment containing straw on the floors could be more eager to investigate this environment, via eating and moving of the straw, when compared to pigs introduced into a pen with no bedding material.

The results from pilot experiments showed that the method used in experiments 1 and 2 should be sufficiently sensitive to detect levels of virus in environmental samples that are similar to levels excreted in urine, faecal or oral samples and in any blood present. However, excretions and secretions from pigs may be spread unevenly in the rooms and detection may require sampling over a larger area than we sampled. Alternative methods for environmental sampling which could be applied to wider areas are likely to improve detection of low levels of contamination by ASFV. One method that could also circumvent safety concerns about collecting infectious materials for analysis used a sponge impregnated with a solvent to inactivate virus and could detect similar amounts of DNA to traditional cotton swabs (Kosowska, Barasona et al. 2021).

In conclusion, the results in this study demonstrate that indirect transmission of ASFV via an environment contaminated with excretions from ASFV-infected pigs is very inefficient when viral DNA levels are similar to those detected in this study and no obvious contamination with blood is present.

## Data availability

The data [values for means and used to build graphs] used to support the findings of this study are available from the corresponding author upon request.

## Acknowledgements

We are very grateful to the staff at IRTA-CReSA and Statens Serum Institut. Especially, we owe great thanks to Guillermo Cantero, Iván Cordon, Joanna Wiacek, María Jesús Navas, Marta Muñoz, Samanta Giler, Xavier Abad and Fie Fisker Brønnum Christiansen for their excellent work.

## Supplementary Materials

**Figure S1:** Detection of ASFV DNA on spiked surfaces using different swabs and elution media. ASFV suspensions (50 µl) with different viral concentrations (10^5^, 10^6^ TCID_50_/ml) were smeared onto petri dishes or dropped on straw and swabbed with either cotton or dust swabs. The swabs and straw were immersed in PBS or culture medium to elute the virus. ASFV DNA was detected by PCR after nucleic acids extraction from the eluate volumes.

**Figure S2:** Detection of infectious ASFV in nasal swabs from inoculated animals in Experiment 2. Porcine bone marrow primary cell cultures were inoculated with nasal swab elution solutions and observed under the light microscope for up to 6 days for development of hemadsorption rosettes (red blood cells that are naturally present in the cultures). Panels are representative of negative (a-b) and positive (c-d) results for the presence of virus. (a) Mock inoculated culture. (b) Nasal swab from Pig 1 at 5 dpi. (c) Nasal swab from Pig 3 at 3 dpi. (d) Nasal swab from Pig 2 at 5 dpi.

